# Use of object detection in camera trap image identification: assessing a method to rapidly and accurately classify human and animal detections for research and application in recreation ecology

**DOI:** 10.1101/2022.01.14.476404

**Authors:** Mitchell Fennell, Christopher Beirne, A. Cole Burton

## Abstract

Camera traps are increasingly used to answer complex ecological questions. However, the rapidly growing number of images collected presents technical challenges. Each image must be classified to extract data, requiring significant labour, and potentially creating an information bottleneck. We applied an object-detection model (MegaDetector) to camera trap data from a study of recreation ecology in British Columbia, Canada. We tested its performance in detecting humans and animals relative to manual image classifications, and assessed efficiency by comparing the time required for manual classification versus a modified workflow integrating object-detection with manual classification. We also evaluated the reliability of using MegaDetector to create an index of human activity for application to the study of recreation impacts to wildlife. In our application, MegaDetector detected human and animal images with 99% and 82% precision, and 95% and 92% recall respectively, at a confidence threshold of 90%. The overall time required to process the dataset was reduced by over 500%, and the manual processing component was reduced by 840%. The index of human detection events from MegaDetector matched the output from manual classification, with a mean 0.45% difference in estimated human detections across site-weeks. Our test of an open-source object-detection model showed it performed well in partially classifying a camera trap dataset, significantly increasing processing efficiency. We suggest that this tool could be integrated into existing camera trap workflows to accelerate research and application by alleviating data bottlenecks, particularly for surveys processing large volumes of human images. We also show how the model and workflow can be used to anonymize human images prior to classification, protecting individual privacy.

**Impact Statement:** We developed and tested a workflow for classifying camera trap images that integrated an existing object-detection model with manual image classification. Our workflow demonstrates an increase in efficiency of 500% over manual labelling, and additionally includes a method to anonymize human images prior to archiving and classification. We provide an example of the application of these tools to ease data processing, particularly for studies focused on recreation ecology which record high volumes of human images. Data lags due to processing delays have the potential to result in sub-optimal conservation decisions, which may be alleviated by accelerated processing. To our knowledge, this is the first in-depth assessment of the practical application of such technology to real world workflows focused on human detections.

## 1. Introduction

In the ongoing quest to better understand and conserve wildlife populations, non-invasive sampling methods have become increasingly important (Zemanova, 2020). One such method is the use of motion-activated remote cameras, or camera traps, which allow researchers to collect extensive observational data while minimally disturbing the wild species of interest, for a relatively low monetary cost (Burton et al., 2015; Caravaggi et al., 2017; Glover-Kapfer et al., 2019; Rowcliffe et al., 2014). Common uses of this technology include species inventories, surveys of occupancy, or calculation of relative abundance indices; however, more recent techniques include estimation of population density (Augustine et al., 2018; Burgar et al., 2018; Jacques et al., 2019; Rich et al., 2014) and analysis of animal behaviour (Caravaggi et al., 2017; Frey et al., 2017). Efforts to standardize camera trap methods and metadata are facilitating cross-project collaboration, paving the way for larger scale syntheses (Forrester et al., 2016; Scotson et al., 2017; Steenweg et al., 2017).

One of the strengths of camera traps is that they can also sample human activity, making simultaneous monitoring of human-wildlife interactions possible. This holds particular promise for studies of recreation ecology, where cameras distributed throughout networks of trails or other recreational corridors provide simultaneous insights on human and wildlife use of habitat in space and time (Baker & Leberg, 2018; George & Crooks, 2006; Kays et al., 2017; Naidoo & Burton, 2020). With improved reliability, and decreasing costs facilitating increased accessibility of camera traps, the number of images collected by researchers continues to grow (Glover-Kapfer et al., 2019; Steenweg et al., 2017). While the use of camera traps provides an excellent framework for multifaceted investigation of fauna worldwide, a major work bottleneck is transforming raw images into usable data for statistical analyses: camera traps often produce high volumes of images, easily reaching terabytes of data.

Once collected by cameras, each image must be reviewed and classified by species, and may be further classified by characteristics of the individual(s) photographed (e.g. age, sex, and behaviour). With many mid-to-large scale projects amassing millions of photos and reaching terabytes of data in less than a year, the time committed to processing these data becomes increasingly unmanageable, ballooning time and monetary budgets. This issue is particularly prominent in recreation ecology studies, where the number of human images captured may frequently be in the millions, with the additional concern of respecting human privacy adding complexity to the issue (Sandbrook et al., 2018, 2021). In some cases, these ethical concerns lead to deletion of all human images, reducing the possibility for detailed assessment of human-wildlife interactions (Naidoo and Burton, 2020). This loss of information on direct human pressures on wildlife is particularly relevant when considering the growing impacts of increasing anthropogenic impacts worldwide (Nickel et al., 2020). Loss of direct management applicability due to a time lag between data collection and reporting is an important disconnect which may result from extensive inefficiencies in processing, limiting an otherwise strong methodology (Merkle et al., 2019; Norouzzadeh et al., 2021). Common strategies to overcome this bottleneck include the use of undergraduate volunteers, contract employees, or community scientists (Lasky et al., 2021; Swanson et al., 2016; Willi et al., 2019). While these strategies assist in accelerating image processing, they do not address the baseline issue that manual processing of images is often extremely labour intensive.

Machine learning methods provide a promising avenue for reducing dependence on manual classification in camera trap projects. Deep learning, a subclass of machine learning, uses artificial neural networks to process information (Lamba et al., 2019). Artificial neural networks are foundationally based on the neural layout seen in biological systems, allowing computer-based algorithms to “learn” based on training data, in order to accurately process similar yet distinct data at a later time. A rapidly expanding subfield of this technology is computer vision, where a multilayered model is trained on a large number of previously classified images, and then applied to new images in order to assign classifications without human interaction (Weinstein, 2018). Researchers have proposed the use of machine learning to identify species in camera trap images (Tabak et al., 2019; Willi et al., 2019; Yu et al., 2013), but tests thus far show performance of these models is generally unsuitable for robust use in real world ecological research (Schneider et al., 2020; Tabak et al., 2019). One of the key current shortcomings is low accuracy when applied to “out-of-pool” samples not seen in training, particularly in new geographical areas without extensive context specific re-training (Schneider et al., 2020).

While future directions to overcome such issues appear promising (e.g. active and transfer learning, Beery et al., 2020; Norouzzadeh et al., 2021), a pragmatic compromise between entirely manual and entirely automated classification of camera trap data is the use of object detection models to assist in filtering images into relevant classes, allowing increased efficiency for manual processing (Beery et al., 2018; Beery, Morris, & Yang, 2019; Greenberg et al., 2019). A key limit to the widespread adoption of such tools by ecologists is a lack of external validation of their performance, slowing adoption of new technologies into practice (Christin et al., 2021).

The release of open-source detection models provides an opportunity for more independent evaluation and application of these tools. One such model, MegaDetector, is a free, openly available, system agnostic object detection model created by Microsoft specifically for the processing of camera trap data (Beery, Morris, & Yang, 2019; Microsoft, 2020). Trained on millions of images from across the world, this model is designed to detect three object classes within images: humans, animals, and vehicles, and can thus be implicitly used to detect images that are blank (i.e. no objects of those classes). Automated processing of images into these classes has the potential to be conducted far faster than could be completed manually by humans and is generally limited only by computer processing speeds (Microsoft, 2020).

While MegaDetector is presented as an effective tool for accelerating data processing, quantification of performance is crucial prior to widespread implementation into real-world workflows (Christin et al., 2021). A key consideration in the evaluation of machine learning model outputs is that these models do not truly “categorize” objects in images, but rather provide a confidence value pertaining to the likelihood of that image containing each object class. While seemingly trivial, it is crucial to establish a threshold against which to measure model performance, as altering confidence thresholds can significantly alter performance measures such as precision, recall, and F-Score. Beyond skewing evaluation of performance, changing thresholds can also change the interpretation of model derived results, potentially leading to different conclusions from the same data.

Here we explore the use of MegaDetector in streamlining camera trap data processing, with a specific focus on projects wishing to quantify large numbers of human images in the context of recreation ecology. Through testing this method on a set of manually classified data, we seek to answer the general question: **can existing object detection models assist in accelerating extraction of ecologically relevant indices from camera trap data?** Specifically, we evaluate the performance of MegaDetector to: *i)* accurately and precisely classify human and wildlife images; ii) produce independent human detection events at a scale commonly used in ecological analysis (site-week), and; iii) increase processing efficiency in comparison to a fully manual workflow. Finally, we also provide an example of direct use of the MegaDetector output to automatically anonymize human images to preserve individual privacy.

## 2. Methods

### 2.1 Manual classification

Images were collected from 36 camera traps in Cathedral Provincial Park, British Columbia, Canada, that were deployed from July 1, 2019 to October 1, 2020 as part of a project investigating the impacts of human recreational activity on wildlife habitat use. Fourteen camera traps were set on human hiking trails, two were on a private road which also serves as a hiking trail, and twenty were set off-trail. Off-trail cameras were deployed in locations a minimum of 300 meters from established trails, resulting in very few human detections other than the research team. Images were classified to the species level, including humans, by a mix of undergraduate volunteers, research assistants, and graduate students. All images were classified manually in a database developed by the UBC WildCo lab. Following initial classification, all classifications were reviewed by the project leader (author MF) to ensure consistency and accuracy across individuals and sites. The average number of images classified per hour was recorded across classifiers to allow comparison to automation assisted approaches. For this analysis, we used 159 272 classified images, of which 74 190 contained humans and 19 120 contained animals, with the remainder being blank images or vehicles. The ratio of human to animal images was 3.88:1.

### 2.2 Object-identifier assisted classification

The same set of images were processed via MegaDetector in order to identify images containing humans, vehicles, and animals, to allow comparison to manual classification outputs.

MegaDetector version 4.1 was run either locally on a desktop (Intel i9-9000 series CPU, 32GB RAM, and an NVIDIA RTX-2080ti GPU), or on a Microsoft Azure NC6s_V3 virtual machine (6 Intel Xeon vCPU’s, 128GB RAM, and an NVIDIA Tesla V100 GPU).

We additionally developed and deployed a human blurring program, which uses the outputs from MegaDetector to obscure individual human identities (see: https://github.com/WildCoLab/WildCo-FaceBlur). This tool uses the output file from MegaDetector, which provides classifications by category, bounding box coordinates around the detection, as well as a confidence value for each detection. Using this information, the blurring program applies a gaussian blur within bounding boxes classified as human above a user defined confidence threshold. Users interact with the program via an R (R Core Team, 2020) interface, which allows specification of a confidence threshold and a level of blurriness, while being familiar for many ecologists. The blurring process itself occurs via Python (Python Software Foundation, 2021) to maximize image handling speed, which is called from the R interface via the package reticulate (Ushey et al., 2021).

Once images were classified manually and processed via MegaDetector, we tested three aspects of object-detector performance to determine efficacy, and also compared the time required to complete processing for each method.

### 2.3 Assessing object-detector classification of human and wildlife images

First, we determined whether images classified manually as containing one or more humans were also classified as “human” by MegaDetector. We summarized classification results as a confusion matrix of true positive (TP), true negative (TN), false positive (FP) and false negative (FN), and used these values to calculate accuracy 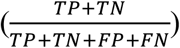, precisio 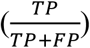, recall 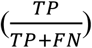, specificity 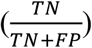 and F-Score 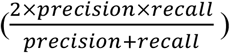. The F-Score is the harmonic mean of the precision and recall, representing a quantification of the tradeoffs between false positives and false negative results, with scores closer to one representing increased model performance. We used a MegaDetector confidence threshold of 90% based on initial sensitivity testing, with a goal of maximizing precision. Second, we applied the same comparison to images containing animals. As MegaDetector does not classify to species beyond detecting an animal, we pooled all animal identifications across species from the manual set for comparison. While the MegaDetector “animal” output requires further manual classification prior to analyses, comparing this class between methods provides broad insight about performance.

### 2.4 Evaluating object-detection based index of human activity

We assessed the reliability of the automated Megadetector classification of human detections as a replacement for manual classification. Camera traps provide a continuous measure of changes in human and animal detections over time, which can be used to analyze temporal trends. We used a time period of 1 week as a relevant index for assessing variation in recreation activity over time. Raw image detections are commonly summarized into independent detection events for statistical analyses of camera trap data, with a minimum time threshold specified between successive images of the same species in order to reduce repeated counting of the same individuals in consecutive triggers and thereby increase independence of observations (Burton et al., 2015). In this case, we used five minutes as the independence threshold between successive events. We grouped the independent events by site-week, resulting in a count of the number of human detection events at each camera trap site for a total of 2052 site-weeks. We compared the count of detection events per site-week from the manual classification to the count of detection events per site-week from MegaDetector (above a 90% confidence threshold) to generate an absolute and percent difference for each site-week, as well as the mean percentage difference across all site weeks. We also calculated a correlation coefficient for the relationship between the two classification methods.

### 2.5 Quantifying gains in efficiency from an object-identifier assisted workflow

To compare efficiency between the manual and automated image classification methods, we recorded the mean number of images processed per hour in our fully manual workflow by having five individuals self-report the time to classify 10 000 images and taking the mean rate across classifiers. We then compared this classification rate to the same metric for our workflow in which MegaDetector was used first, followed by manual classification to species of all animal images. As animal images still require manual classification following detection via MegaDetector, we calculated the time required for the full dataset to be processed on our local computer with an NVIDIA RTX 2080ti GPU and added the time to classify the 19 120 animal images manually at the average manual rate to provide an estimated time for the full workflow.

## 3. Results

### 3.1 Classification of human images

MegaDetector detected human images with 97.2% accuracy when compared to the manual classification (Table 1a), showing that 94.7% of images containing a human were correctly assigned to this class at the 90% confidence threshold. Precision for human image detection was 0.99, recall was 0.95, and specificity was 0.99. The F-Score was 0.97, and the misclassification rate was 2.8%. The correlation coefficient between manually identified and MegaDetector identified human images was 0.96 (Fig 1a). More error was observed at camera sites with more images, suggesting a consistent rate of error, with MegaDetector being more likely to undercount the number of human images at the 90% confidence threshold (Fig 1a). A precision-recall curve for confidence thresholds from 0-0.9 is presented in supplemental Figure S1.

**Figure 1.**
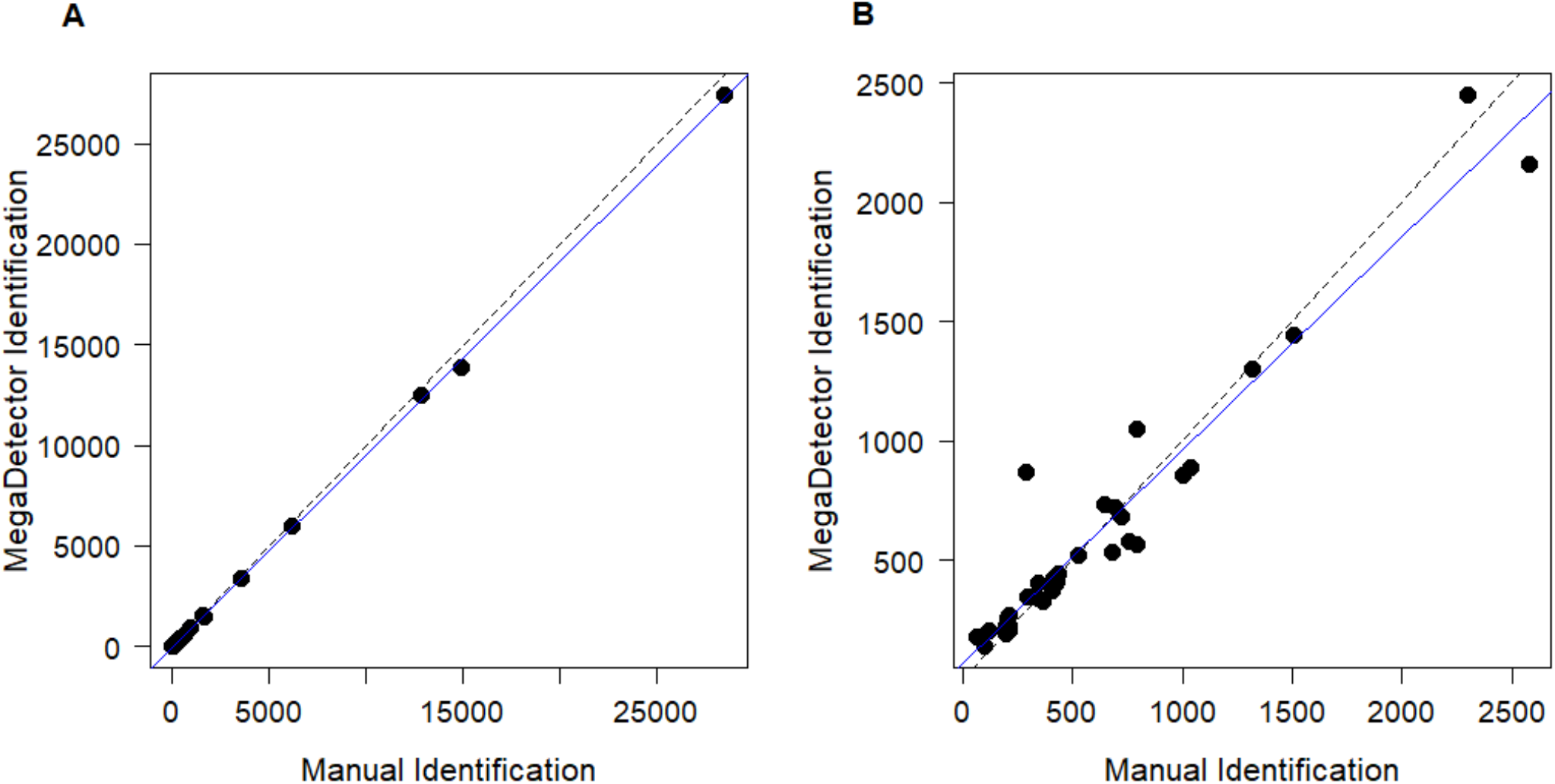
(A) Human image detections at each remote camera station. The horizontal axis is the number of manually identified human images at each site, and the vertical axis is the number of MegaDetector classified human images at each site with a 90% confidence threshold. The dashed line is a 1:1 regression, and the solid blue line is the regression between manual and automated classification at each site (correlation coefficient 0.96). (B) Animal image detections at each remote camera station. The horizontal axis is the number of manually identified animal images at each site, and the vertical axis is the number of MegaDetector classified animal images at each site with a 90% confidence threshold. The dashed line is a 1:1 regression, and the solid blue line is the regression between manual and automated classification at each site (correlation coefficient 0.89).

**Table 1.** (A) Confusion matrix of object-identifier classification of human images, above a 90% confidence threshold. The percentage in the top left represents the true positive rate, top right represents the false positive rate, bottom left is the false negative rate, and bottom right is the true negative rate. (B) Confusion matrix of object-identifier classification of animal images, above a 90% confidence threshold. The percentage top left represents the true positive rate, top right represents the false positive rate, bottom left is the false negative rate, and bottom right is the true negative rate.

|  |  | Manual |  |
| --- | --- | --- | --- |
|  |  | Human | No Human |
| MegaDetector | Human | 70275<br>(94.7%) | 617<br>(0.7%) |
|  | No Human | 3915<br>(5.3%) | 84465<br>(99.3%) |

B
|  |  | Manual |  |
| --- | --- | --- | --- |
|  |  | Animal | No Animal |
| MegaDetector | Animal | 17654<br>(92.3%) | 4005<br>(2.9%) |
|  | No Animal | 1466<br>(7.7%) | 136125<br>(97.1%) |

### 3.2 Classification of animal images

Animal image detection by MegaDetector had an accuracy of 96.6% at the 90% confidence threshold (Table 1b). Table 1b shows 92.3% of all animal images were correctly identified by MegaDetector at this threshold. The precision for object-identifier based animal image classification was 0.82, the sensitivity 0.92, and the specificity 0.97. The F-Score was 0.87 and the misclassification rate 3.4%. The correlation coefficient for camera site level animal image classification was 0.89 with increased variation per site when compared to human images (Fig 1b). MegaDetector was once again slightly conservative, more commonly underestimating the number of true animal images at the 90% confidence threshold.

### 3.3 Independent human events

Object-identifier assisted classification of human image events was highly correlated at the site-week level, showing strong correspondence between manual and MegaDetector based inference (Fig. 2). The mean number of independent human events per site-week as determined by manual classification was 4.45, with a range from 0 to 207 events. The mean difference in human events identified by MegaDetector was −0.0024 events per site week, with a range from −3 to 4. The mean percentage difference in the number of independent human detection events across 2052 site-weeks was 0.45%. The correlation coefficient of the relationship between object-detection and manual classification of independent human detection events was 0.99.

**Figure 2.**
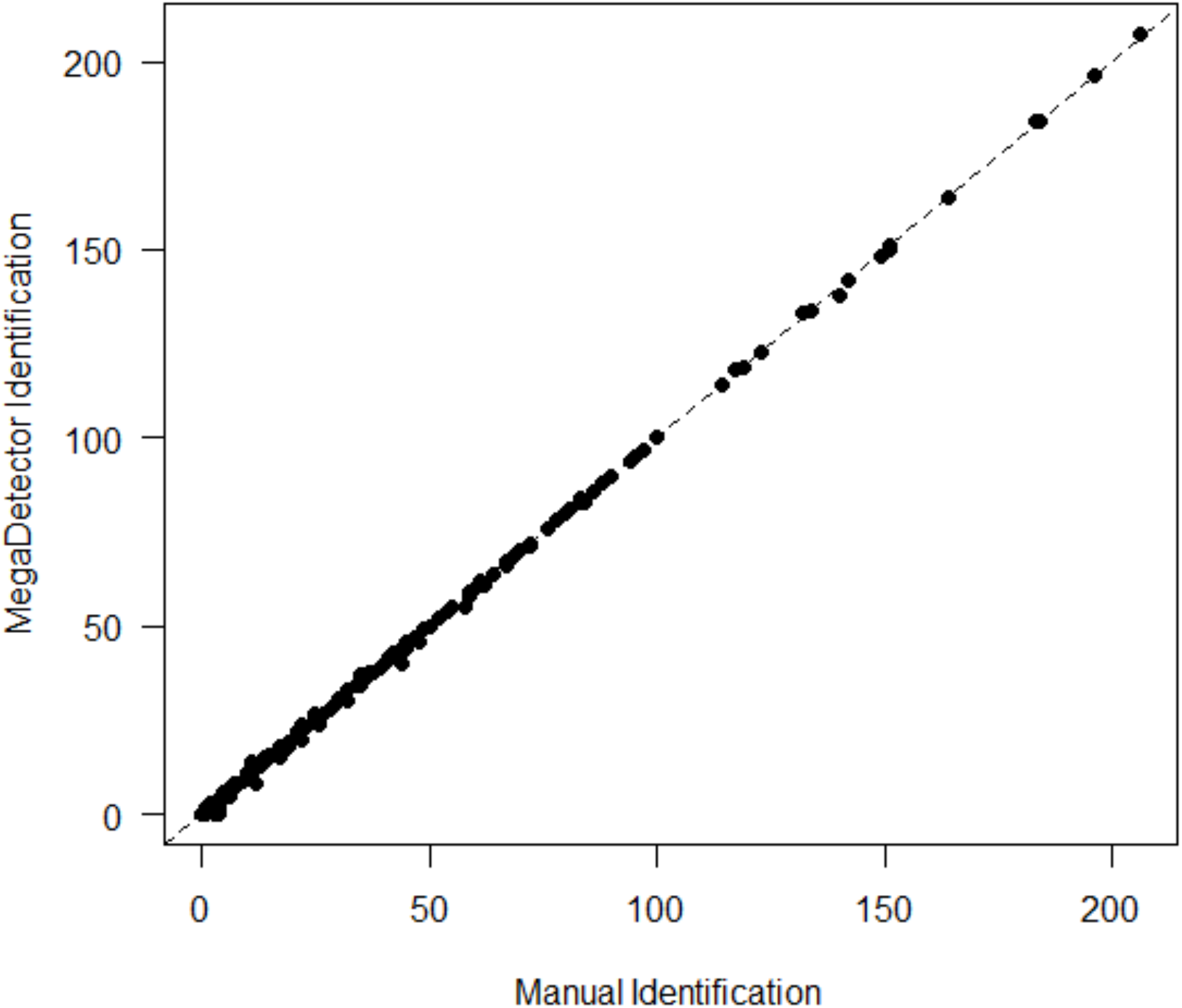
Independent human detection events across 2052 camera trap site-weeks as manually classified vs. classified via object-detection, using a 90% confidence threshold. The dashed horizontal line is a 1:1 relationship.

### 3.4 Time benchmarks

For human classifiers, classification speed ranged from 300 to 1000 images per hour, with experienced classifiers being faster than novice classifiers. The mean rate across five classifiers was 500 images per hour.

MegaDetector inference speed is highly hardware dependent, largely related to GPU throughput. Our local machine with an NVIDIA RTX 2080ti GPU processed images at an average of rate of 1.8 images per second (6480 images/hour), while our virtual machine with an NVIDIA Tesla V100 GPU processed images at a rate of 2.3 images per second (8280 images/hour). Preliminary testing subsequent to these analyses with an NVIDIA RTX 3090 GPU produced classification rates of 2.8 images per second (10 080 images/hour).

Using the mean manual processing rate of 500 images per hour, our dataset took an estimated 319 person-hours to classify 159 272 images. The time to process the entire dataset via MegaDetector with our slowest computer was 25 hours, with an additional 38 hours to manually classify the animal images, for a total of 63 hours for the MegaDetector based workflow. The MegaDetector based workflow was 506% faster than the entirely manual workflow, with 40% of the total time not requiring human input or supervision, resulting in an 840% decrease in the manual processing time.

### 3.5 Human-blurring

We deployed our human-blurring tool on all images prior to manual classification to protect human privacy of recreationists recorded on camera traps. As this tool is dependent on MegaDetector outputs, our success at blurring these data was the same as those reported above (Fig 1a, Table 1a). By providing an optional parameter for the level of blur applied to each human classified bounding box, we retained the ability to identify humans within images during manual classification while preventing individual identification, preserving privacy. This process required very little manual input beyond specifying the file path to the folder containing all images, as well as an output directory.

## 4. Discussion

This study provides an example of the effective application of an existing open source object-detection model to greatly accelerate the classification of camera trap image data. This will increase the efficiency of processing large volumes of human detections, which will be particularly beneficial in studies quantifying effects of human activity on wildlife, such as in recreation ecology. Where timely data is essential in supporting conservation decision making, extensive time lags between collection of data and useable recommendations can exacerbate existing disconnects between research and effective action (Dubois et al., 2020; Habel et al., 2013; Sands, 2012). This acceleration in processing may assist in narrowing gaps between researchers and managers by providing timely information to support decision making (Cvitanovic et al., 2016; Lemieux et al., 2018; Merkle et al., 2019).

Using an object-detection model assisted workflow, we achieved increases in processing speed of over 500% while maintaining high accuracy and precision of image classification for a realistic ecological dataset. By implementing an existing tool which is openly available (Microsoft MegaDetector; Beery, Morris & Yang, 2019; Microsoft, 2020), we were able to adjust an existing workflow to integrate machine learning for human image detection—a task on which current algorithms perform strongly--while continuing manual classification of animal species identification, for which machine learning performance is currently weaker. While the field of computer vision is rapidly improving the ability for automated species identification (Beery et al., 2020; Gomez Villa et al., 2017), existing models are not commonly generalizable to new environments or species, resulting in performance which is not yet acceptable for broad application by ecologists without significant error checking and correction (Beery, Morris, Yang, et al., 2019; Glover-Kapfer et al., 2019; Schneider et al., 2020). The workflow tested here provides significant increases in processing efficiency while keeping the “human in the loop” for a quality comparable to fully manual camera trap image processing, with minimal correction needed, and without requiring extensive model retraining for a geographic region unseen in training.

These gains in efficiency are particularly relevant in the context of recreation ecology, where the number of human images may vastly exceed the number of animal images. Recreation presents a unique and understudied pressure to wildlife, with species and geographically specific responses to different activities varying (Baker & Leberg, 2018; Boyle & Samson, 1985; Kays et al., 2017; Naidoo & Burton, 2020; Nickel et al., 2020). One contributing factor to this knowledge gap is the difficulty in quantifying recreation pressures, particularly while simultaneously measuring wildlife (Balmford et al., 2015). Camera traps provide an excellent opportunity to overcome this issue, though large numbers of human images may be overwhelming for researchers to process in an effective manner. In the case of our study, the ratio of human to animal images was 3.88, with our object-identifier assisted workflow resulting in an 840% reduction in manual processing hours. In cases where this ratio is higher, we predict that the increase in processing efficiency will be even greater.

In addition to the improvement in processing efficiency, we demonstrated that MegaDetector facilitates the blurring of human images prior to data processing. The human ethics of camera trapping has received attention recently (Sandbrook et al., 2021; Sharma et al., 2020) and the ability to anonymize images before they are viewed by human observers or archived is in our opinion considered best practice. To this end, we provide a script (https://github.com/WildCoLab/WildCo-FaceBlur) which takes the human-labelled objects identified by MegaDetector and blurs them using Python via a simple R interface, which is familiar for many ecologists. Important to note is that despite the high performance of MegaDetector shown here, some human images were missed by the object-detection process, which may preclude obscuring every human image captured by camera traps. In our anecdotal experience, the vast majority of miss-labelled human images were cases of a single foot, hand, or backpack moving through the edge of the image field-of-view, presenting very limited ability for individual identification whether blurred or not.

Potential limitations of using object detection methods are the perceived technical knowledge required and the need for high-performance computing hardware. Regarding the former, MegaDetector is accompanied by detailed yet simple instructions to assist practitioners in applying the model to their own data (Microsoft, 2020). In terms of hardware infrastructure, these models will run on nearly any modern computer, though inference speed is significantly increased with the use of a dedicated performance GPU. We suggest two options for access to such hardware: purchasing or upgrading a computer to use locally, or the use of a virtual machine such as those hosted by Amazon AWS, Microsoft Azure, Google Cloud, or various academic institutions. While upfront costs may be high for a suitable computer or GPU, it is relevant to consider these costs in comparison with the labour wages needed to cover the increased processing time for entirely manual classification, particularly as applied across multiple projects over the life of the hardware. Virtual machines (portions of large, high performing computers allocated and accessed remotely) are an accessible option for short-term projects which may not warrant the upfront hardware expense, or to initially trial the efficacy of integrating these tools with your own workflow. Though this option provides the ability to have on demand access to high-performance computing, costs can quickly increase and exceed those of purchasing physical hardware (Tuia et al., 2021).

The promise of camera traps to provide rapid information for applied management of human-wildlife landscapes is currently limited by the rapid increase in data collected worldwide (Ahumada et al., 2020; Steenweg et al., 2017). While this issue has presented a potential limitation to widespread adaptation of this technology, rapid developments in processing technology, such as those in the field of artificial intelligence provide exciting solutions (Ahumada et al., 2020; Glover-Kapfer et al., 2019). Here we provide an example of integrating such tools into an existing workflow in a recreation ecology context, showing the high performance of an openly available model on real data, and suggesting application of such technology by other ecological practitioners. While the future holds further drastic developments for the applicability of these technologies to camera trap data and research, we propose that there is no time like the present for ecologists to use all available tools to accelerate their research, particularly in situations where alleviating data lags may facilitate effective conservation decisions.

## Acknowledgements

We thank all team members involved in data collection and image processing, including Jamie Clark, Jamie Fennell, Lucas Friesen, Isla Francis, Eve Hurd, Taylor Justason, Joey Krahn, Jessica Low, Michael Procko, Debra Sinarta, and Charles White. We thank the staff of BC Parks for their support, particularly Melanie Percy, James Quayle, Kirk Safford, and Kevin Wilson. We also sincerely thank the Microsoft AI for Earth team, particularly Dan Morris and Siyu Yang for their dedicated work on MegaDetector, as well as generous support with implementation, answering a few too many emails, and for comments from DM which greatly improved this manuscript.

## Author Contributions

Conceptualization: M.F; C.B; A.C.B. Methodology: M.F; C.B; A.C.B. Data curation: M.F; A.C.B. Data Analysis: M.F; C.B. Writing original draft: M.F. All authors approved the final submitted draft and provided substantial editing to the final draft.

## Data Availability

Data availability: The summary data and code that support the findings of this study are openly available on Github at https://github.com/mitch-fen/FennellBeirneBurton_2022. Raw image data is not made available due to privacy constraints.

## Ethical Standards

Data was collected, processed, and stored as approved under University of British Columbia Animal Care Permit A18-0234, and Human Ethics Permit H21-01424.

## Supplemental Material

**Figure S1.**
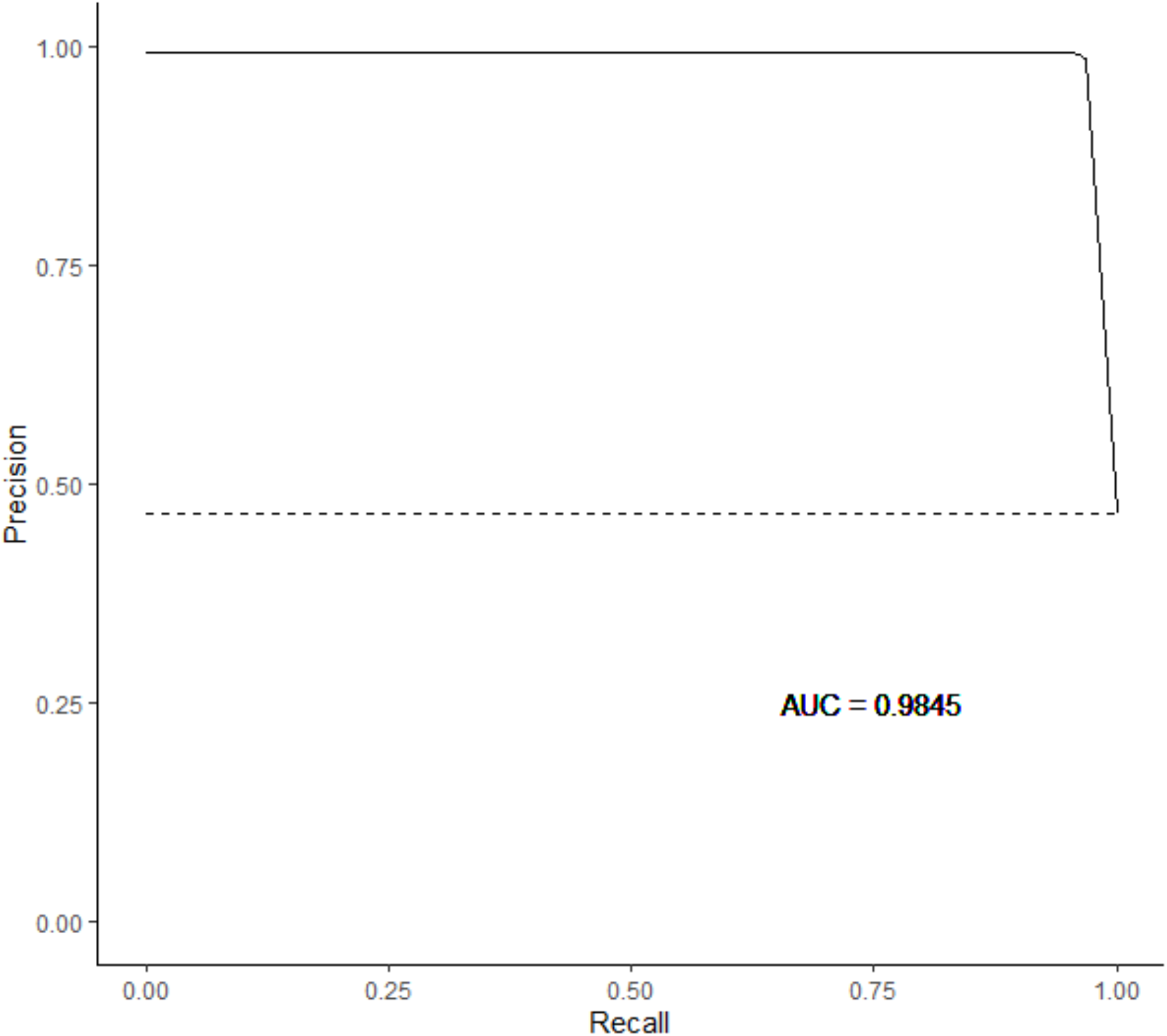
Precision-recall curve for human image detection by MegaDetector, for 9 confidence threshold values from 0 to 0.9. The AUC value is the area under the precision-recall curve, as calculated using the trapezoidal rule.

## References

Ahumada, J. A., Fegraus, E., Birch, T., Flores, N., Kays, R., O’Brien, T. G., Palmer, J., Schuttler, S., Zhao, J. Y., Jetz, W., Kinnaird, M., Kulkarni, S., Lyet, A., Thau, D., Duong, M., Oliver, R., & Dancer, A. (2020). Wildlife Insights: A Platform to Maximize the Potential of Camera Trap and Other Passive Sensor Wildlife Data for the Planet. Environmental Conservation, 47(1), 1–6. https://doi.org/10.1017/S0376892919000298

Augustine, B. C., Royle, J. A., Kelly, M. J., Satter, C. B., Alonso, R. S., Boydston, E. E., & Crooks, K. R. (2018). Spatial capture–recapture with partial identity: An application to camera traps. The Annals of Applied Statistics, 12(1), 67–95. https://doi.org/10.1214/17-AOAS1091

Baker, A. D., & Leberg, P. L. (2018). Impacts of human recreation on carnivores in protected areas. PLOS ONE, 13(4), e0195436. https://doi.org/10.1371/journal.pone.0195436

Balmford, A., Green, J. M. H., Anderson, M., Beresford, J., Huang, C., Naidoo, R., Walpole, M., & Manica, A. (2015). Walk on the Wild Side: Estimating the Global Magnitude of Visits to Protected Areas. PLOS Biology, 13(2), e1002074. https://doi.org/10.1371/journal.pbio.1002074

Beery, S., Morris, D., & Yang, S. (2019). Efficient Pipeline for Camera Trap Image Review. ArXiv:1907.06772 [Cs]. http://arxiv.org/abs/1907.06772

Beery, S., Morris, D., Yang, S., Simon, M., Norouzzadeh, A., & Joshi, N. (2019). Efficient Pipeline for Automating Species ID in new Camera Trap Projects. Biodiversity Information Science and Standards, 3, e37222. https://doi.org/10.3897/biss.3.37222

Beery, S., van Horn, G., & Perona, P. (2018). Recognition in Terra Incognita. ArXiv:1807.04975 [Cs, q-Bio]. http://arxiv.org/abs/1807.04975

Beery, S., Wu, G., Rathod, V., Votel, R., & Huang, J. (2020). Context R-CNN: Long Term Temporal Context for Per-Camera Object Detection. ArXiv:1912.03538 [Cs, Eess, q-Bio]. http://arxiv.org/abs/1912.03538

Boyle, S. A., & Samson, F. B. (1985). Effects of Nonconsumptive Recreation on Wildlife: A Review. Wildlife Society Bulletin, 13(2), 8.

Burgar, J. M., Stewart, F. E. C., Volpe, J. P., Fisher, J. T., & Burton, A. C. (2018). Estimating density for species conservation: Comparing camera trap spatial count models to genetic spatial capture-recapture models. Global Ecology and Conservation, 15, e00411. https://doi.org/10.1016/j.gecco.2018.e00411

Burton, A., Neilson, E., Moreira, D., Ladle, A., Steenweg, R., Fisher, J., Bayne, E., & Boutin, S. (2015). Wildlife camera trapping: A review and recommendations for linking surveys to ecological processes. JOURNAL OF APPLIED ECOLOGY, 52(3), 675–685. https://doi.org/10.1111/1365-2664.12432

Caravaggi, A., Banks, P. B., Burton, A. C., Finlay, C. M. V., Haswell, P. M., Hayward, M. W., Rowcliffe, M. J., Wood, M. D., Pettorelli, N., & Sollmann, R. (2017). A review of camera trapping for conservation behaviour research. Remote Sensing in Ecology and Conservation, 3(3), 109–122. https://doi.org/10.1002/rse2.48

Christin, S., Hervet, É., & Lecomte, N. (2021). Going further with model verification and deep learning. Methods in Ecology and Evolution, 12(1), 130–134. https://doi.org/10.1111/2041-210X.13494

Cvitanovic, C., McDonald, J., & Hobday, A. J. (2016). From science to action: Principles for undertaking environmental research that enables knowledge exchange and evidence-based decision-making. Journal of Environmental Management, 183, 864–874. https://doi.org/10.1016/j.jenvman.2016.09.038

Dubois, N. S., Gomez, A., Carlson, S., & Russell, D. (2020). Bridging the research-implementation gap requires engagement from practitioners. Conservation Science and Practice, 2(1). https://doi.org/10.1111/csp2.134

Forrester, T., O’Brien, T., Fegraus, E., Jansen, P. A., Palmer, J., Kays, R., Ahumada, J., Stern, B., & McShea, W. (2016). An Open Standard for Camera Trap Data. Biodiversity Data Journal, 4(4), e10197–8. https://doi.org/10.3897/BDJ.4.e10197

Frey, S., Fisher, J. T., Burton, A. C., & Volpe, J. P. (2017). Investigating animal activity patterns and temporal niche partitioning using camera-trap data: Challenges and opportunities. Remote Sensing in Ecology and Conservation, 3(3), 123–132. https://doi.org/10.1002/rse2.60

George, S. L., & Crooks, K. R. (2006). Recreation and large mammal activity in an urban nature reserve. Biological Conservation, 133(1), 107–117. https://doi.org/10.1016/j.biocon.2006.05.024

Glover-Kapfer, P., Soto-Navarro, C. A., Wearn, O. R., Rowcliffe, M., & Sollmann, R. (2019). Camera-trapping version 3.0: Current constraints and future priorities for development. Remote Sensing in Ecology and Conservation, 5(3), 209–223. https://doi.org/10.1002/rse2.106

Gomez Villa, A., Salazar, A., & Vargas, F. (2017). Towards automatic wild animal monitoring: Identification of animal species in camera-trap images using very deep convolutional neural networks. Ecological Informatics, 41, 24–32. https://doi.org/10.1016/j.ecoinf.2017.07.004

Greenberg, S., Godin, T., & Whittington, J. (2019). Design patterns for wildlife-related camera trap image analysis. Ecology and Evolution, 9(24), 13706–13730. https://doi.org/10.1002/ece3.5767

Habel, J. C., Gossner, M. M., Meyer, S. T., Eggermont, H., Lens, L., Dengler, J., & Weisser, W. W. (2013). Mind the gaps when using science to address conservation concerns. Biodiversity and Conservation, 22(10), 2413–2427. https://doi.org/10.1007/s10531-013-0536-y

Jacques, C. N., Klaver, R. W., Swearingen, T. C., Davis, E. D., Anderson, C. R., Jenks, J. A., Deperno, C. S., & Bluett, R. D. (2019). Estimating density and detection of bobcats in fragmented midwestern landscapes using spatial capture–recapture data from camera traps. Wildlife Society Bulletin, 43(2), 256–264. https://doi.org/10.1002/wsb.968

Kays, R., Parsons, A. W., Baker, M. C., Kalies, E. L., Forrester, T., Costello, R., Rota, C. T., Millspaugh, J. J., McShea, W. J., & Toit, J. (2017). Does hunting or hiking affect wildlife communities in protected areas? Journal of Applied Ecology, 54(1), 242–252. https://doi.org/10.1111/1365-2664.12700

Lamba, A., Cassey, P., Segaran, R. R., & Koh, L. P. (2019). Deep learning for environmental conservation. Current Biology, 29(19), R977–R982. https://doi.org/10.1016/j.cub.2019.08.016

Lasky, M., Parsons, A., Schuttler, S., Mash, A., Larson, L., Norton, B., Pease, B., Boone, H., Gatens, L., & Kays, R. (2021). Candid Critters: Challenges and Solutions in a Large-Scale Citizen Science Camera Trap Project. Citizen Science: Theory and Practice, 6(1), 4. https://doi.org/10.5334/cstp.343

Lemieux, C. J., Groulx, M. W., Bocking, S., & Beechey, T. J. (2018). Evidence-based decision-making in Canada’s protected areas organizations: Implications for management effectiveness. FACETS, 3(1), 392–414. https://doi.org/10.1139/facets-2017-0107

Merkle, J. A., Anderson, N. J., Baxley, D. L., Chopp, M., Gigliotti, L. C., Gude, J. A., Harms, T. M., Johnson, H. E., Merrill, E. H., Mitchell, M. S., Mong, T. W., Nelson, J., Norton, A. S., Sheriff, M. J., Tomasik, E., & Vanbeek, K. R. (2019). A collaborative approach to bridging the gap between wildlife managers and researchers. The Journal of Wildlife Management, 83(8), 1644–1651. https://doi.org/10.1002/jwmg.21759

Microsoft. (2020). AI for Earth camera trap image processing API. (4.1) [Computer software]. Microsoft. https://github.com/microsoft/CameraTraps/blob/master/megadetector.md

Naidoo, R., & Burton, A. C. (2020). Relative effects of recreational activities on a temperate terrestrial wildlife assemblage. Conservation Science and Practice, 2(10). https://doi.org/10.1111/csp2.271

Nickel, B. A., Suraci, J. P., Allen, M. L., & Wilmers, C. C. (2020). Human presence and human footprint have non-equivalent effects on wildlife spatiotemporal habitat use. Biological Conservation, 241(Journal Article), 108383. https://doi.org/10.1016/j.biocon.2019.108383

Norouzzadeh, M. S., Morris, D., Beery, S., Joshi, N., Jojic, N., & Clune, J. (2021). A deep active learning system for species identification and counting in camera trap images. Methods in Ecology and Evolution, 12(1), 150–161. https://doi.org/10.1111/2041-210X.13504

Python Software Foundation. (2021). Python 3 (3.7) [Computer software]. http://www.python.org/

R Core Team. (2020). R: A language and environment for statistical computing. R Foundation for Statistical Computing. https://www.R-project.org/

Rich, L. N., Kelly, M. J., Sollmann, R., Noss, A. J., Maffei, L., Arispe, R. L., Paviolo, A., De Angelo, C. D., Di Blanco, Y. E., & Di Bitetti, M. S. (2014). Comparing capture-recapture, mark-resight, and spatial mark-resight models for estimating puma densities via camera traps. Journal of Mammalogy, 95(2), 382–391. https://doi.org/10.1644/13-MAMM-A-126

Rowcliffe, J. M., Kays, R., Kranstauber, B., Carbone, C., & Jansen, P. A. (2014). Quantifying levels of animal activity using camera trap data. Methods in Ecology and Evolution, 5(11), 1170–1179. https://doi.org/10.1111/2041-210X.12278

Sandbrook, C., Clark, D., Toivonen, T., Simlai, T., O’Donnell, S., Cobbe, J., & Adams, W. (2021). Principles for the socially responsible use of conservation monitoring technology and data. Conservation Science and Practice, 3(5). https://doi.org/10.1111/csp2.374

Sandbrook, C., Luque-Lora, R., & Adams, W. M. (2018). Human Bycatch: Conservation Surveillance and the Social Implications of Camera Traps. Conservation and Society, 16(4), 493–504. https://doi.org/10.4103/cs.cs_17_165

Sands, J. P. (2012). Wildlife science: Connecting research with management. CRC Press. https://doi.org/10.1201/b12139

Schneider, S., Greenberg, S., Taylor, G. W., & Kremer, S. C. (2020). Three critical factors affecting automated image species recognition performance for camera traps. Ecology and Evolution, 10(7), 3503–3517. https://doi.org/10.1002/ece3.6147

Scotson, L., Johnston, L. R., Iannarilli, F., Wearn, O. R., Mohd-Azlan, J., Wong, W. M., Gray, T. N. E., Dinata, Y., Suzuki, A., Willard, C. E., Frechette, J., Loken, B., Steinmetz, R., Moßbrucker, A. M., Clements, G. R., & Fieberg, J. (2017). Best practices and software for the management and sharing of camera trap data for small and large scales studies. Remote Sensing in Ecology and Conservation, 3(3), 158–172. https://doi.org/10.1002/rse2.54

Sharma, K., Fiechter, M., George, T., Young, J., Alexander, J. S., Bijoor, A., Suryawanshi, K., & Mishra, C. (2020). Conservation and people: Towards an ethical code of conduct for the use of camera traps in wildlife research. Ecological Solutions and Evidence, 1(2). https://doi.org/10.1002/2688-8319.12033

Steenweg, R., Hebblewhite, M., Kays, R., Ahumada, J., Fisher, J. T., Burton, C., Townsend, S. E., Carbone, C., Rowcliffe, J. M., Whittington, J., Brodie, J., Royle, J. A., Switalski, A., Clevenger, A. P., Heim, N., & Rich, L. N. (2017). Scaling-up camera traps: Monitoring the planet’s biodiversity with networks of remote sensors. Frontiers in Ecology and the Environment, 15(1), 26–34. https://doi.org/10.1002/fee.1448

Swanson, A., Kosmala, M., Lintott, C., & Packer, C. (2016). A generalized approach for producing, quantifying, and validating citizen science data from wildlife images: Citizen Science Data Quality. Conservation Biology, 30(3), 520–531. https://doi.org/10.1111/cobi.12695

Tabak, M. A., Norouzzadeh, M. S., Wolfson, D. W., Sweeney, S. J., Vercauteren, K. C., Snow, N. P., Halseth, J. M., Di Salvo, P. A., Lewis, J. S., White, M. D., Teton, B., Beasley, J. C., Schlichting, P. E., Boughton, R. K., Wight, B., Newkirk, E. S., Ivan, J. S., Odell, E. A., Brook, R. K.,… Miller, R. S. (2019). Machine learning to classify animal species in camera trap images: Applications in ecology. Methods in Ecology and Evolution, 10(4), 585–590. https://doi.org/10.1111/2041-210X.13120

Tuia, D., Kellenberger, B., Beery, S., Costelloe, B. R., Zuffi, S., Risse, B., Mathis, A., Mathis, M. W., van Langevelde, F., Burghardt, T., Kays, R., Klinck, H., Wikelski, M., Couzin, I. D., van Horn, G., Crofoot, M. C., Stewart, C. V., & Berger-Wolf, T. (2021). Seeing biodiversity: Perspectives in machine learning for wildlife conservation. ArXiv:2110.12951 [Cs]. http://arxiv.org/abs/2110.12951

Ushey, K., Allaire, J., & Tang, Y. (2021). reticulate: Interface to Python (1.19) [Computer software]. https://rstudio.github.io/reticulate/

Weinstein, B. G. (2018). A computer vision for animal ecology. Journal of Animal Ecology, 87(3), 533–545. https://doi.org/10.1111/1365-2656.12780

Willi, M., Pitman, R. T., Cardoso, A. W., Locke, C., Swanson, A., Boyer, A., Veldthuis, M., Fortson, L., & Gaggiotti, O. (2019). Identifying animal species in camera trap images using deep learning and citizen science. Methods in Ecology and Evolution, 10(1), 80–91. https://doi.org/10.1111/2041-210X.13099

Yu, X., Wang, J., Kays, R., Jansen, P. A., Wang, T., & Huang, T. (2013). Automated identification of animal species in camera trap images. EURASIP Journal on Image and Video Processing, 2013(1), 52. https://doi.org/10.1186/1687-5281-2013-52

Zemanova, M. A. (2020). Towards more compassionate wildlife research through the 3Rs principles: Moving from invasive to non-invasive methods. Wildlife Biology, 2020(1). https://doi.org/10.2981/wlb.00607

